# A microcircuit model for initiation and reward-dependent termination of hippocampal replay

**DOI:** 10.64898/2026.01.18.700234

**Authors:** Hugo Musset, Tomoki Fukai

## Abstract

Sharp-wave ripples (SWRs) serve as a key network state in which the hippocampus replays spatiotemporal patterns that reflect animals’ experiences. However, how the hippocampal circuitry controls the generation, duration, and termination of these events remains unclear. To clarify how behaviorally salient locations are encoded into replay, we constructed a biologically plausible model of area CA3, including three classes of inhibitory neurons — parvalbumin-positive basket cells, cholecystokinin-positive basket cells (CCKBC), and theta-OFF, ripple-ON cells (TORO). In our model, replay and SWRs emerge from TORO-mediated release of CCKBC inhibition onto pyramidal cells, as hypothesized by experiments. Furthermore, our model suggests that short-term depression at TORO-CCKBC synapses controls replay duration and identifies a mechanism through which replay overrepresents reward-associated locations. Our model predicts that replays terminating at these locations are shorter, which we confirm in experimental data. Overall, this study provides a mechanistic framework for SWRs and replay dynamics in the hippocampus.

## Introduction

Decades of experimental and theoretical evidence have identified hippocampal sharp-wave ripples (SWR) — short high-frequency (140-200 Hz) oscillatory bouts occurring during slow-wave sleep and behavioral immobility — as a substrate for memory consolidation and retrieval (Wilson & McNaughton 1994; Buzsáki 2015). Disrupting and prolonging these SWRs impairs and enhances memory performance, respectively (Girardeau *et al*. 2009; Jadhav *et al*. 2012; Fernández-Ruiz *et al*. 2019).

Nested within SWRs, replay describes trajectory through space by sequentially reactivating place cells and provides a framework for synaptic plasticity (Nádasdy *et al*. 1999; Carr *et al*. 2011; Sadowski *et al*. 2016; Ó lafsdóttir *et al*. 2018). Replay can be either forward or reverse, and that ratio differs for replay occurring during slow-wave-sleep or quiet wakefulness (Foster & Wilson 2006; Diba & Buzsáki 2007; Csicsvari *et al*. 2007). Replay is known to overrepresent salient locations such as landmarks and rewards, hinting at its importance in goal-driven behavior (Pfeiffer & Foster 2013; Igata *et al*. 2021).

Generation of SWRs is thought to be the result of interactions between the pyramidal cell (PC) population and a class of recurrently connected parvalbumin-expressing basket cells (PVBC, Schlingloff *et al*. 2014; Stark *et al*. 2014). In contrast, little is known about SWR termination, with spike-frequency adaptation and neuronal fatigue proposed as explanatory mechanisms (English *et al*. 2014; Buzsáki 2015). Intriguingly, SWR duration follows a long-tailed distribution reflecting memory demands (Fernández-Ruiz *et al*. 2019), suggesting the existence of a content-aware termination mechanism that is actively mediated (English *et al*. 2014). Recent findings have revealed new cellular-level actors intertwined with SWRs: a population of cholecystokinin-expressing basket cells (CCKBC, Dudok *et al*. 2021) with activity patterns opposite to PVBC and PC, and a population of inhibitory neurons active exclusively during SWRs, receiving direct PC input and projecting strongly to local interneurons (TORO – theta-off, ripple-on, Szabo *et al*. 2022), making them ideal candidates for controlling the SWR microcircuitry.

To elucidate how the hippocampal circuitry controls the generation, duration, and termination of replay events and how their end points represent behaviorally salient locations, we built a biologically plausible microcircuit model of replay and SWRs in the strongly recurrent CA3 hippocampal area (Sammons, Vezir, *et al*. 2024) comprising PC, PVBC, CCKBC, and TORO. We found that SWRs occur spontaneously through TORO-mediated release of CCKBC inhibition. The associated replays are driven by the recurrent projection between place cells reflecting the relative position of their place fields and terminate through stochastic changes in CCKBC activity. Unexpectedly, TORO-to-CCKBC inhibition enables our model to generate a wide range of SWR durations, such as those observed in experiments. Furthermore, we implemented a simple mechanistic encoding of reward position in our network at the PC-CCKBC synapses, resulting in a termination mechanism modulated by the content of replay. Previously, phenomenological state-space models of memory and replay assumed replay termination without explicitly modeling the underlying network mechanisms (Mattar & Daw 2018; Hunt *et al*. 2021; Bakermans *et al*. 2025). Our biologically plausible network model reproduced the experimentally observed overrepresentation of reward (Pfeiffer & Foster 2013; Widloski & Foster 2022) along with a novel observation that replays terminating at reward are shorter than those terminating elsewhere, which we verified in experimental data.

Altogether, these results provide a biology-constrained mechanism for the generation and termination of replay and SWRs with a wide range of possible durations in the context of goal-directed learning. Our model suggests that TORO cells play pivotal roles in initiating and prolonging, but not directly terminating, these events.

## Results

To explore the mechanisms of SWR generation and termination and how they are affected by replay content, we built a microcircuit model of rodent CA3 hippocampal area comprising pyramidal cells, two interneuron classes linked to SWRs (PVBC and CCKBC), and TORO cells. The number of cells has been chosen to match that of a 400-*µ*m-thick slice, experimentally determined to be the minimum size required for spontaneous SWR generation *in vitro* (Hájos *et al*. 2013; Schlingloff *et al*. 2014). The relative abundance of PV and CCK interneurons and TORO cells is also well characterized (Jinno & Kosaka 2010; Whissell *et al*. 2015; Szabo *et al*. 2022) and can be scaled down to the mimicked slice size, yielding 8000 PCs, 150 PVBCs, and 160 CCKBCs and 50 TORO cells. All cells were modeled as conductance-based adaptive exponential integrate-and-fire point-process neurons (AEIF) and connected uniformly according to data derived from anatomical studies when available with values fixed within a biologically plausible range otherwise. Figure 1A shows the general connectivity scheme. To reflect the dense pyramidal CA3 connectivity, the modeled PC-PC weight matrix (Fig. 1B) is strongly recurrent and follows a noisy distance-dependent scheme. Place cells (50% of PCs) were assigned a place field in a virtual 3m-long periodic track. Thus, cells with adjacent place fields are more likely to be strongly connected than cells farther apart/non-place-coding-cells. We optimized the values of parameters, including the strengths of synaptic connections, such that our network can replicate biologically realistic properties of SWRs (see Materials and Methods). The resulting PC-PC weights follow a heavy-tailed distribution (Fig. 1C), associated experimentally and computationally with a wide range of neuronal and network properties, including but not limited to SWR generation and propagation (Omura *et al*. 2015; Ecker *et al*. 2022; Buzsáki & Mizuseki 2014). The network receives feedforward granule cell (GC) -like uncorrelated Poisson spike trains onto PC and CCKBC (Acsády *et al*. 1998), driving the activity of pyramidal cells around 0.1Hz. This low-level baseline activity effectively reproduces quiet wakefulness/slow-wave sleep behavior as opposed to theta states characterized by strong rhythmic input from the medial septum (Borhegyi *et al*. 2004). Single-cell and network parameters/connectivity are detailed in the Materials and Methods section.

**Figure 1:**
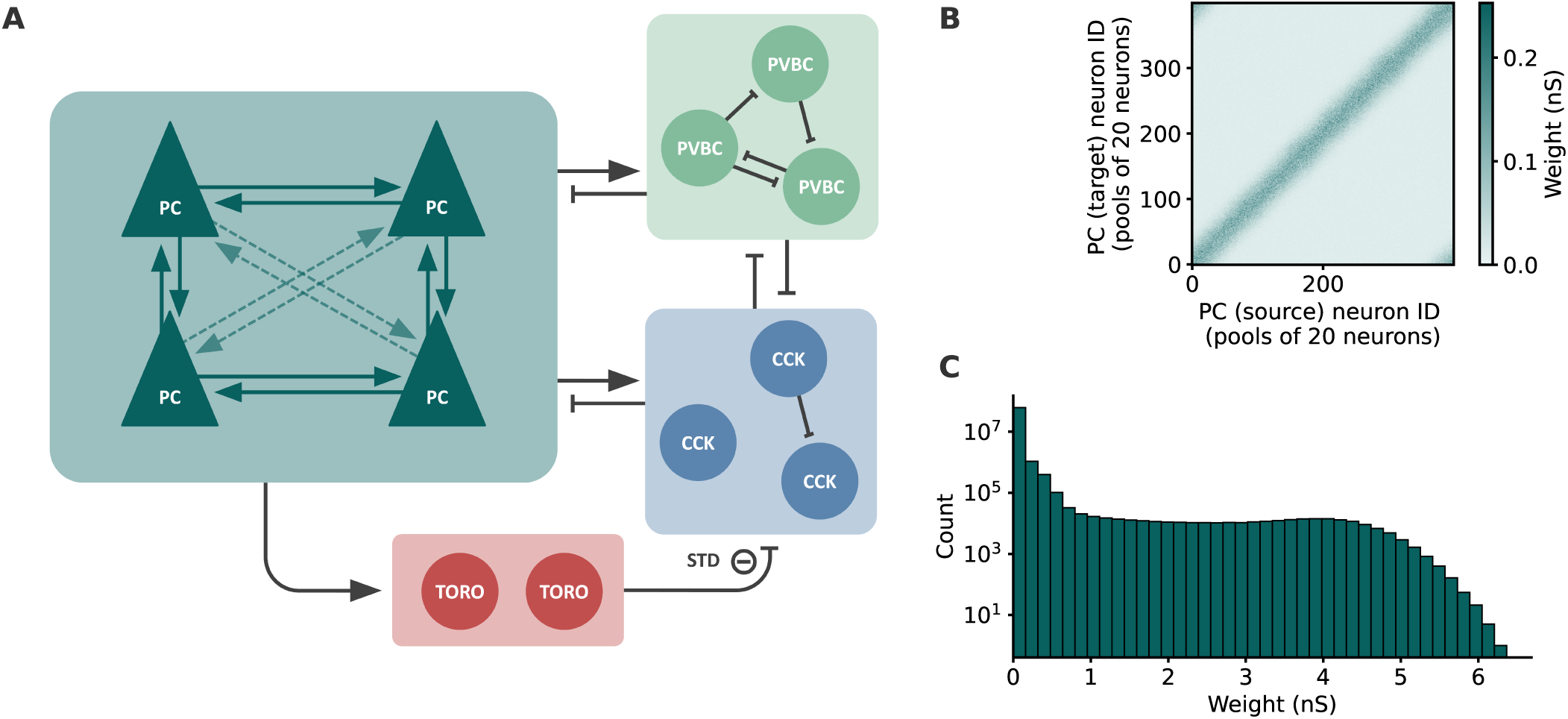
Network structure. ***A***, Network connectivity scheme. ***B*** , Recurrent connectivity matrix between pyramidal cells (PC), pooled in groups of 20 cells for visibility. Cells are ordered according to the center of their place field on the periodic environment. Place and non-place cells have alternating indices. ***C*** , weight distribution from the PC-PC matrix in B.

### Replay is spontaneously generated

Our model replicates the various features of spontaneous hippocampal activity observed in experiments. Activity in the PC population is stochastically amplified and can trigger network-wide high-activity transient events (Fig. 2A). Such events reproduce many aspects of experimentally-recorded SWRs such as their frequency (Fig. 2 B) and local field potential (LFP) profile (Fig. 2C). As was suggested experimentally, the accompanying ripple-frequency oscillation (150-250 Hz, Buzsáki 2015) is generated through the interplay of the PC and PVBC populations and relies on the strong recurrent connectivity in the PVBC network (Schlingloff *et al*. 2014). Throughout this study, these events will be referred to as sharp-wave ripples. A shift in PC inhibitory tone from CCKBC to PVBC characterizes alternating baseline and SWR states, respectively (Dudok *et al*. 2021). PC activity throughout the SWR reflects the relative position of place cells and is shaped by the network’s recurrent connectivity. We can decode place information from the temporal pattern of this activity, known as replay, to reconstruct a coherent trajectory through the environment in which the cells have a place field (Fig. 2D, see Materials and Methods). We note that our model maintains the realistic firing rates observed in excitatory and inhibitory cells during SWRs (Fig. 2E) (Lasztóczi *et al*. 2011; Varga *et al*. 2014; Dudok *et al*. 2021; Szabo *et al*. 2022).

**Figure 2:**
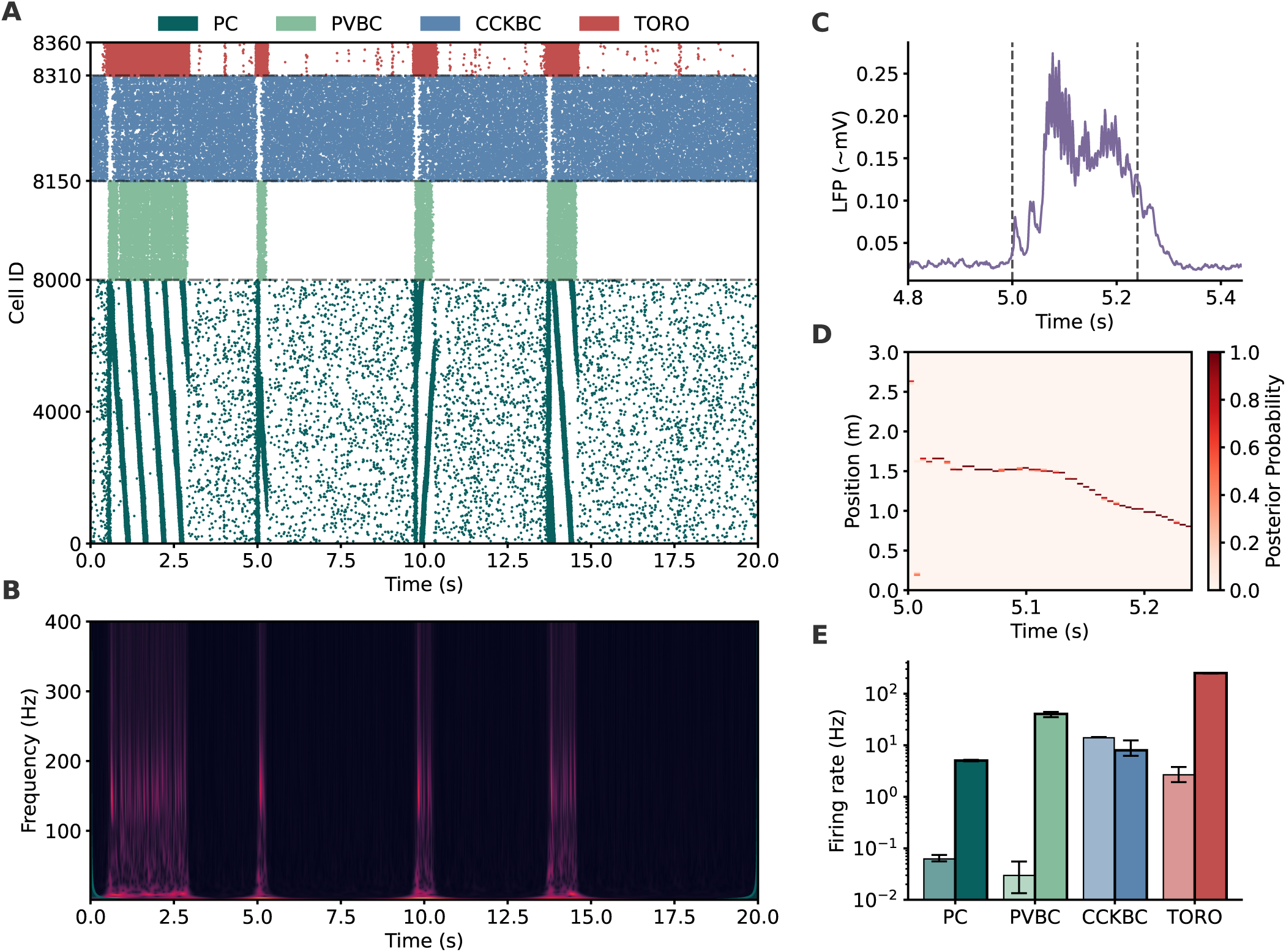
Spontaneous generation of SWRs and replay. ***A***, Network raster with all 8000 PCs, 150 PVBCs, 160 CCKBCs, and 50 TORO cells. PCs ordered by their place field in the periodic environment. ***B*** , Associated scaleogram obtained from the estimated LFP. ***C*** , Estimated LFP around the time of a SWR in ***A***. ***D*** , Decoded replay from a SWR in ***A***. ***E*** , Population firing rates for all populations in the network, sorted by rates outside of SWRs (light tones, left) and rates within SWRs (dark tones, right). Error bars mark the 25th and 75th percentiles from 5000 s of simulation.

### SWR generation through TORO-mediated disinhibition

In order to expose the population-level activity patterns leading to SWR generation and alternation of PVBC-CCKBC inhibitory tone, we selectively excited or inhibited single populations in our model. These manipulations of neural population activity reveal the division of labor among the different inhibitory neuron subtypes in generating SWRs. In our model, the generation of SWRs requires appropriate switching among the activities of PVBC, CCKBC, and TORO.

First of all, the excitation of a spatially coherent subset of PCs is generally sufficient for SWR generation (Fig. 3A1). Inhibition of CCKBC either through TORO activation (Fig. 3A2) or direct inhibition (Fig. 3A3) leads to the disinhibition of the ripple network consisting of PC and PVBC, thus triggering a SWR. In contrast, the activation of PVBC (such that their activity level matches that observed during SWRs — e.g., population-level rhythmic firing at the ripple frequency) fails to activate PC and hence is insufficient for SWR generation. This result is consistent with that of the FINO (fast inhibitory neuronal oscillations) model of SWRs proposed by Schlingloff *et al*. (2014), in which excitatory activity must first build up in the PC population before being amplified: Otherwise, the PC population is too strongly inhibited for any activity to emerge. The initial non-specific increase in PC activity in Fig. 3A2-3 reflects increased excitability of PC due to the lower overall inhibitory tone.

**Figure 3:**
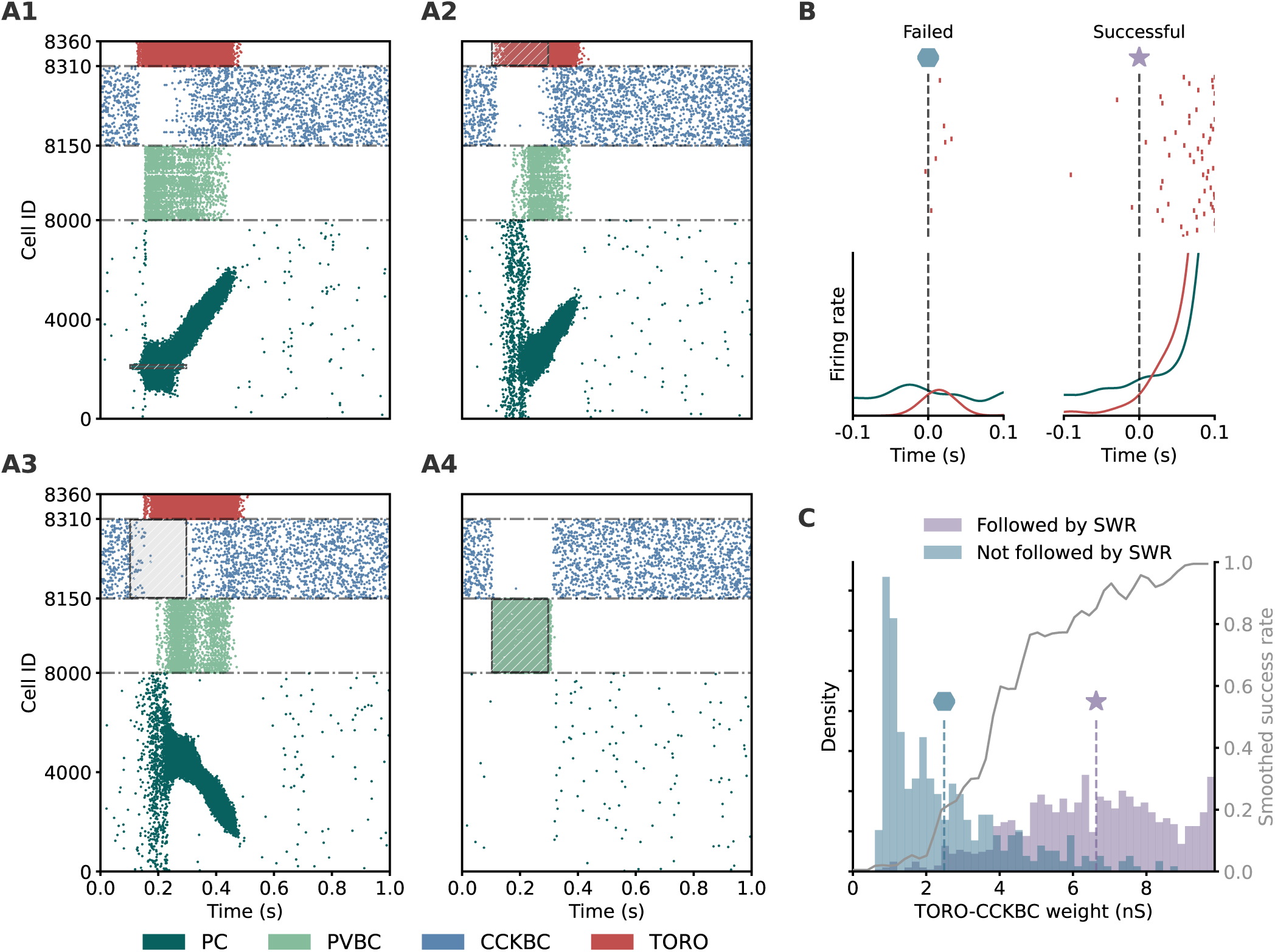
Evoked SWRs and requirement for successful initiation. ***A***, Network rasters following external stimulation (excitation or inhibition). ***A*1**, Excitation of a subset of 100 PCs with adjacent place fields for 200 ms (shaded box). ***A*2**, Excitation of TORO cells for 200 ms (shaded box). ***A*3**, Inhibition of CCKBCs for 200 ms (shaded box). ***A*4**, Excitation of PVBCs for 200 ms (shaded box). ***B*** , TORO spike trains around a failed (left) and successful (right) SWR initiation, with the corresponding TORO and PC firing rates below. Time is relative to when TORO firing rate crosses a certain threshold (see Materials and Methods for details). ***C*** , Average synaptic weight from TORO to CCKBC for many failed and successful spontaneous SWR initiation attempts. Colored dashed lines mark the mean weight for each group. Grey curve: SWR initiation success rate obtained by summing successful and failed events (smoothed with 0.2 nS gaussian kernel).

In sum, our model predicts that the activation of TORO disinhibits the PC-PVBC ripple network by inhibiting CCKBC on top of the dynamics of the PVBC-CCKBC reciprocal network proposed by Evangelista *et al*. (2020), where a pro- and an anti-SWR population compete and inhibit each other. This allows PC activity to be amplified according to the recurrent connectivity reflecting spatial proximity between place cells (winner-take-all amplification), leading to SWR and sequence replay. By rapidly responding to changes in PC activity, TORO effectively flips the bistability switch between PVBC and CCKBC-dominated inhibition.

### TORO-CCKBC STD determines successful SWR initiation

We next analyzed how the short-term dynamics at the TORO-CCKBC synapses affect SWR generation. Compared to their PVBC counterparts, TORO cells respond to fluctuations in PC activity with a greater sensitivity, and high activity epochs in the TORO population tend to be accompanied by SWRs in the model (Fig. 2A). However, outside SWRs, there are brief epochs of TORO activity that no SWRs follow immediately. We refer to these increases in TORO firing rate as SWR initiation attempts and categorize them into successful (leading to a SWR: Fig. 3B, right) and unsuccessful (not immediately followed by a SWR: Fig. 3B, left). We asked what distinguishes successful attempts from unsuccessful attempts. Initially, PC and TORO firing rate trajectories are similar in both attempts, suggesting that TORO fails to release CCKBC inhibition at the later stage. Indeed, plotting the instantaneous weight values of the TORO to CCKBC synapses around the initiation of these attempts reveals the synaptic weight dependency: SWR attempts are unlikely to be successful when the synaptic weights have not been fully recovered from STD and remain low, whereas the success rate of an attempt increases drastically when the synapses are already recovered (Fig. 3C). These results show that weak TORO-CCKBC synaptic strength in the failed attempt prevented TORO from releasing CCKBC inhibition, stopping further amplification of PC activity that would have led to a SWR. It is noted that the dependency on TORO to CCKBC STD could modify the statistics of SWR generation by posing a refractory period after a SWR event. Experimentally this period is ∼200 ms in slice preparations (Schlingloff *et al*. 2014), although we do not tune our model to reproduce it.

These results make TORO a key actor for initiating SWRs in the CA3 microcircuitry, by selecting PVBC or CCKBC as the main inhibitory source onto PCs and controlling the timescale of their activation.

### CCKBC activity destabilizes SWR events

The duration of SWRs is highly variable and reflects hippocampal engagement and memory demand, such as familiar vs novel environments and memory vs non-memory tasks Fernández-Ruiz *et al*. 2019; Vancura *et al*. 2023; Quigley *et al*. 2023). Such large variability cannot be solely accounted for by a buildup of adaptive currents or a reduction in synaptic drive. It is more likely that an active biological mechanism mediates the termination (English *et al*. 2014; Buzsáki 2015). In our model, SWR termination is consistently associated with an increase in CCKBC firing rate (Fig. 4A). This increase in CCKBC activity can be attributed to decreases in TORO-CCKBC synaptic efficacy, because STD suppresses it after repeated high-frequency TORO firing during SWRs (Fig. 4B).

**Figure 4:**
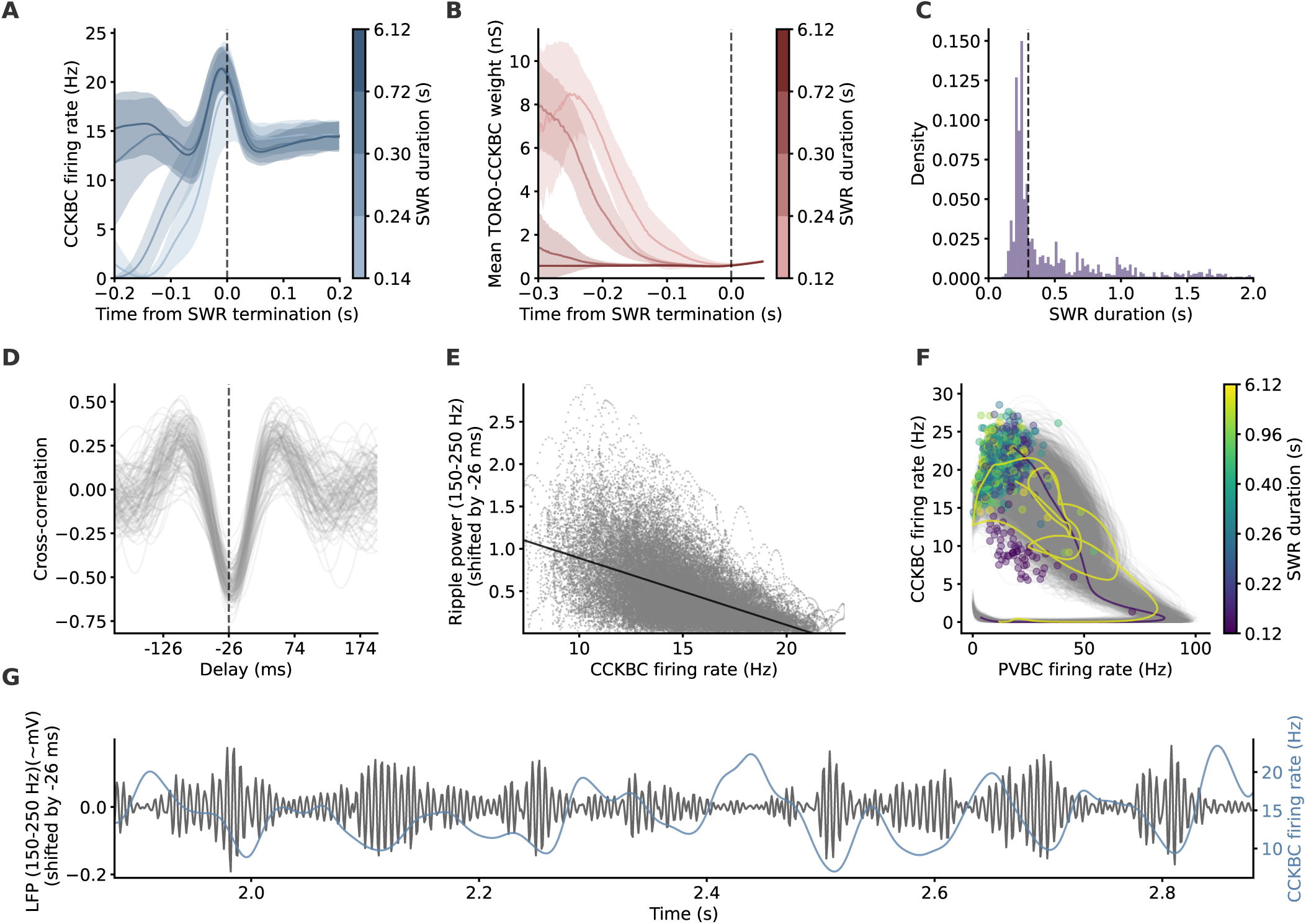
Termination of SWRs by CCKBC. ***A***, Firing rate of CCKBCs around the time of SWR termination, grouped by duration of SWR. Group identity corresponds to consecutive quartiles. ***B*** , TORO-CCKBC synaptic weight around the time of SWR termination, grouped by duration of SWR. ***C*** , Distribution of SWR durations. Dashed line marks the median SWR duration. ***D*** , Cross-correlation between CCKBC firing rate and ripple-frequency-range-band-pass-filtered LFP envelope during SWRs. Each gray line represents one SWR. Dashed line marks the median delay maximizing negative cross-correlation. ***E*** , Ripple-frequency-range-band-pass-filtered LFP envelope shifted by the delay in ***D*** vs. CCKBC firing rate during SWRs. Dark line is best fit line (*y* = −0.079*x* + 1.68). ***F*** , CCKBC and PVBC phase-space trajectories of SWRs. Dots mark the position at termination, and are color-coded by SWR duration. Two example trajectories are shown and are color-coded according to their duration. ***G***, CCKBC firing rate and ripple-frequency-range-band-pass-filtered LFP shifted by the delay in ***D*** during one example SWR from Fig. 2A). Data in ***A***-***B*** and ***D*** -***F*** from 100 50-s long simulations.

Importantly, SWR duration follows a long-tailed distribution (Fig. 4C) in accordance with experimental observations (Fernández-Ruiz *et al*. 2019). In long-duration SWRs in particular, CCKBC activity is already high and TORO-CCKBC weight is already very weak hundreds of ms before the actual termination of the SWR (Figs. 2A and 4A-B). This suggests the possibility that an elevated CCKBC activity is not sufficient to immediately terminate SWRs. SWRs may instead represent an unstable network state, terminated through stochastic changes in the balance between pro (mainly PVBC and TORO) and anti (CCKBC) -SWR activities. We note that largely variable SWR duration is unlikely in a previous model without TORO cells, in which SWRs terminate solely due to short-term depression dynamics at PVBC-to-CCKBC synapses (Evangelista *et al*. 2020).

Under this assumption, we characterized the relationship between CCKBC activity and the ripple oscillation. We found that changes in CCKBC firing rate were consistently followed by opposite changes in ripple amplitude and power (Fig. 4E, G), with a delay of ∼26 ms (Fig. 4D). The phase space of PVBC-CCKBC firing rate reveals that SWRs go through alternating PVBC and CCKBC-dominated phases before CCKBC activity stochastically drives the network strongly enough out of the unstable equilibrium, terminating the SWR (Fig. 4F).

In sum, the balance in pro- and anti-SWR activity governs the time course of SWRs: CCKBC activity increases through synaptic fatigue at the TORO-CCKBC synapses, followed by decreases in the ripple power and the firing rate of the PC population, which further decreases CCKBC activity and strengthens the ripple, providing more excitation to CCKBC, forming a repeating cycle until CCKBC activity eventually terminates the ongoing SWR.

### The role of CCKBC in terminating versus non-terminating SWRs

In the parameter space of synaptic connection strengths, only a small subset of combinations leads to networks that can exhibit SWRs. An even smaller fraction of these networks generate SWRs that can terminate. As such, we explored the robustness of our results against variations in parameter values.

CCKBC receives synaptic inputs from all other neuron groups and is located at the core of our model. It receives feedforward and recurrent excitation from GC and PC, respectively, and inhibition from PVBC and TORO, and excitation-inhibition balance on CCKBC determines the network’s ability to generate SWRs and their properties. We found that when the overall excitation onto CCKBC is too strong, the network is unable to generate SWRs, as strong CCKBC activity prevents PC-mediated TORO recruitment (Fig. 5A1, purple region). In contrast, when the recurrent (SWR-driven) excitation is too weak, CCKBC activity remains low during the SWR and is unable to terminate it (Fig. 5A1, green region). In this region, as the PC-CCKBC weight decreases, the average duration of SWRs increases until a point where SWRs do not terminate within the simulation time window. An example non-terminating SWR is given in Fig. 5A2). We emphasize that a sweet spot where PC-CCKBC excitation is weak outside but strong within SWRs occupies a sufficiently wide region in the parameter space, allowing the model to generate and terminate SWRs in a finite time (Fig. 5A1, blue region).

**Figure 5:**
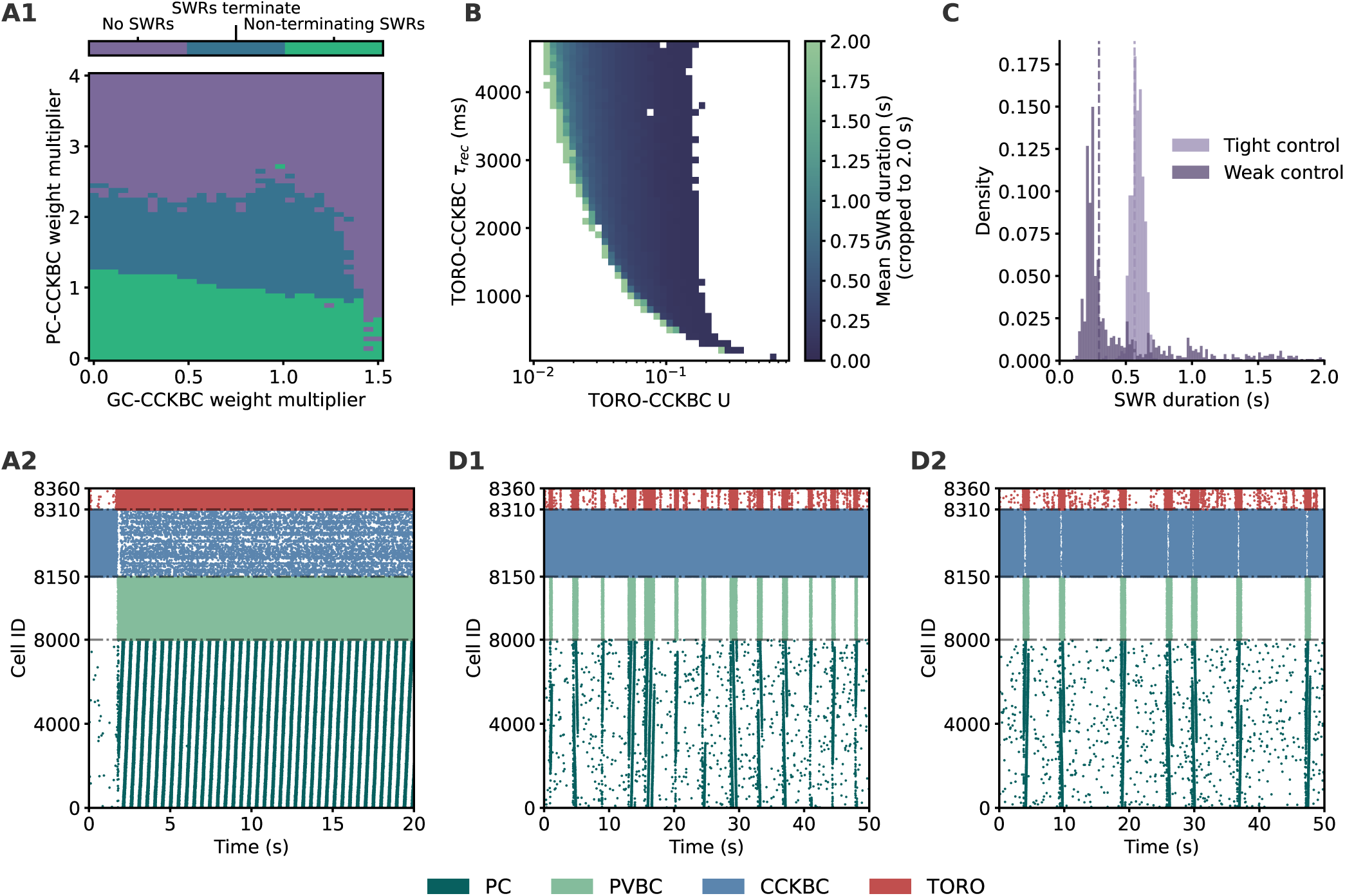
Terminating SWRs are only a subset of SWRs. ***A1*** , Types of SWRs across excitatory inputs to CCKBC (GC and PC). ***A2*** , Example non-terminating SWR. ***B*** , Duration of SWRs across TORO-CCKBC short-term plasticity parameters (recovery time constant and synaptic efficacy). White areas are regions with either no SWRs (right) or non-terminating SWRs (left). For visualization purposes, SWRs are cropped to 2 s maximum duration. ***C*** , Distribution of durations of SWRs produced by the networks in ***D1*** and ***D2*** . ***D1*** , Example network with a weak control on SWR duration. ***D2*** , Example network with a tight control on SWR duration. Each pixel in ***A1*** and ***B*** corresponds to the average of three 50 s-long simulations. Data in ***C*** from 500 50-s long simulations.

### TORO-CCKBC synaptic transmission controls SWR duration

As shown earlier, the time course of SWRs is modulated by the short-term plasticity at the TORO-CCKBC synapses. Indeed, controlling SWR duration is a novel and important feature of our microcircuit model with TORO cells. We systematically analyzed SWR duration throughout the parameter space of short-term depression (the recovery time constant and synaptic efficacy, Tsodyks & Markram 1997) at these synapses to clarify how their dynamics affect the ability of the network to modulate SWR duration. In this analysis, we reduced the parameter space to a subset that can generate SWRs that terminate (Materials and Methods).

Long-duration SWRs form a contiguous region along an increasing synaptic efficacy and a decreasing recovery time constant. Namely, a long SWR can occur when the recovery time constant is large and synaptic efficacy is low, and vice-versa (Fig. 5B). SWRs to the left of this wavefront do not terminate: TORO-CCKBC synapses are weakly depressing, and CCKBC activity remains too low to terminate the SWR. In this region, a persistent SWR becomes the steady state of the network. In contrast, no SWRs are able to appear to the right of the wavefront. TORO-CCKBC synapses are strongly depressing, and the release of CCKBC inhibition is too transient for SWR generation.

Importantly, our network can produce both long-tailed and Gaussian-distributed SWR durations, depending on the choice of network parameters (Fig. 5C). By changing the TORO-CCKBC synaptic parameters from *U* = 0.09 and *τ_rec_* = 700 ms to *U* = 0.02 and *τ_rec_* = 4400 ms(“long-tailed or weak control” and “Gaussian-like or tight control”, respectively), we were able to constrain our model to produce weakly regulated SWRs with various durations or very strongly regulated SWRs with a fixed duration (Fig. 5D1 and D2, respectively).

### Reward-driven replay termination by CCKBC

An intriguing characteristic of hippocampal replay activity is that it overrepresents reward location at termination (Pfeiffer & Foster 2013; Widloski & Foster 2022). How exactly this preference for particular locations can happen remains unclear. This preference is likely crucial for learning a spatial map, as it informs the animal of the locations of reward delivery by confining replay activity within behaviorally meaningful spatial segments. To our knowledge, no previous models have explored a biologically plausible mechanism of this important characteristic of replay activity.

Our previous results showed that CCKBC activity promotes SWR termination in the hippocampal microcircuit. Therefore, we hypothesize that increasing CCKBC activity around a rewarded location increases the likelihood of termination there. A simple yet effective mechanism to achieve this is a reward-distance-dependent modulation of synaptic weights from PC to CCKBC. Namely, we hypothesized that PC-to-CCKBC synapses are stronger around the reward location (Fig. 6B). This modulation may arise from a reward-modulated long-term plasticity rule applied during exploration. However, exploring the explicit mechanism of this modification is beyond the scope of this study. We will discuss the possible mechanisms later.

**Figure 6:**
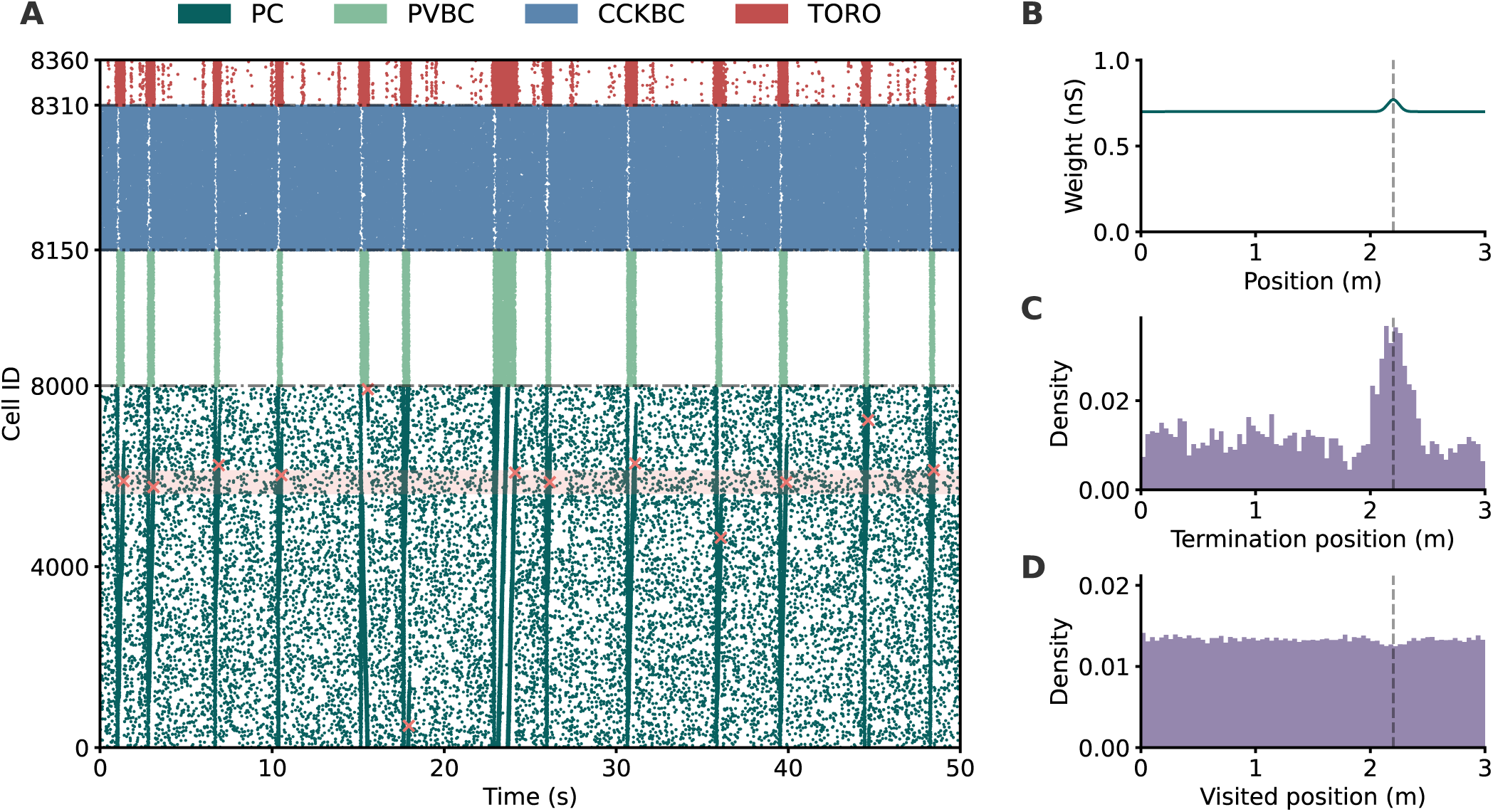
Overrepresentation of reward at termination. ***A***, Network raster. Orange crosses mark SWR termination location and orange shaded area marks reward position +/−10 cm. ***B*** , PC-CCKBC synaptic weight, ordered by center of place field. Dashed line marks reward position. ***C*** , Termination sites of replay. ***D*** , Visited sites during replay. For visualization purposes, the reward-associated modifications in PC-CCKBC synaptic weight used in ***A*** are amplified. Data in ***C*** and ***D*** from 500 50-s long simulations.

The outcome of reward-distance-dependent PC-to-CCKBC synapses is an increased representation of the reward through the termination of replay activity (Fig. 6A, C). Importantly, this increased termination at the reward location is not a consequence of the increased representation of this location during replay. To see this, we calculated the distribution of all spatial points visited by repeated replay events. The obtained distribution shows no signature of the overrepresentation of reward location during replay (Fig. 6D), confirming the above claim. The result, however, may conflict with some experimental observations that demonstrated a more frequent visit by replay to the reward location (Pfeiffer & Foster 2013). This discrepancy may arise from the fact that replay activity progresses randomly in our model, whereas, in reality, it may trace the spatial paths that likely lead to a reward location.

Thus, our model can produce reward overrepresentation through replay termination, a hallmark of goal-directed learning, and further explores the paradigm introduced by Jahnke *et al*. (2015) and expanded upon by Ecker *et al*. (2022), mechanistically linking SWR dynamics and actual replay content. Our results describe a simple biologically plausible mechanism through which replay content directly affects the ongoing SWR, terminating it at behaviorally relevant locations.

### Context-specific reward-terminating replays are shorter

The mechanism described above predicts a strong CCKBC activation around reward-associated areas. Any replay trying to pass through this area is more likely to terminate there, as shown above. As such, it is expected that replay activities that terminate at reward locations are shorter than their counterparts that terminate elsewhere: if they didn’t go through a reward area, they would not be exposed to higher CCKBC firing rates and likely terminate “naturally” later at another random location.

We first verified that CCKBC activity during replay is indeed higher near the reward location. To this end, we simulated a linear environment with solid boundaries at either end and a reward located at 2.2 m along the track, and collected many replay episodes in this environment from our network modified as described in Fig. 6B. By averaging the CCKBC activity for each spatially decoded position over repeated runs, we observed a bump of activity around the reward location (Fig. 7A), confirming the relationship of an elevated CCKBC activity with reward. Secondly, we sorted replay events by the position at which they terminated (within 10 cm of the reward or elsewhere). As hypothesized, this analysis revealed that replays terminating at or near the reward are shorter than those terminating elsewhere in our model (Fig. 7B).

**Figure 7:**
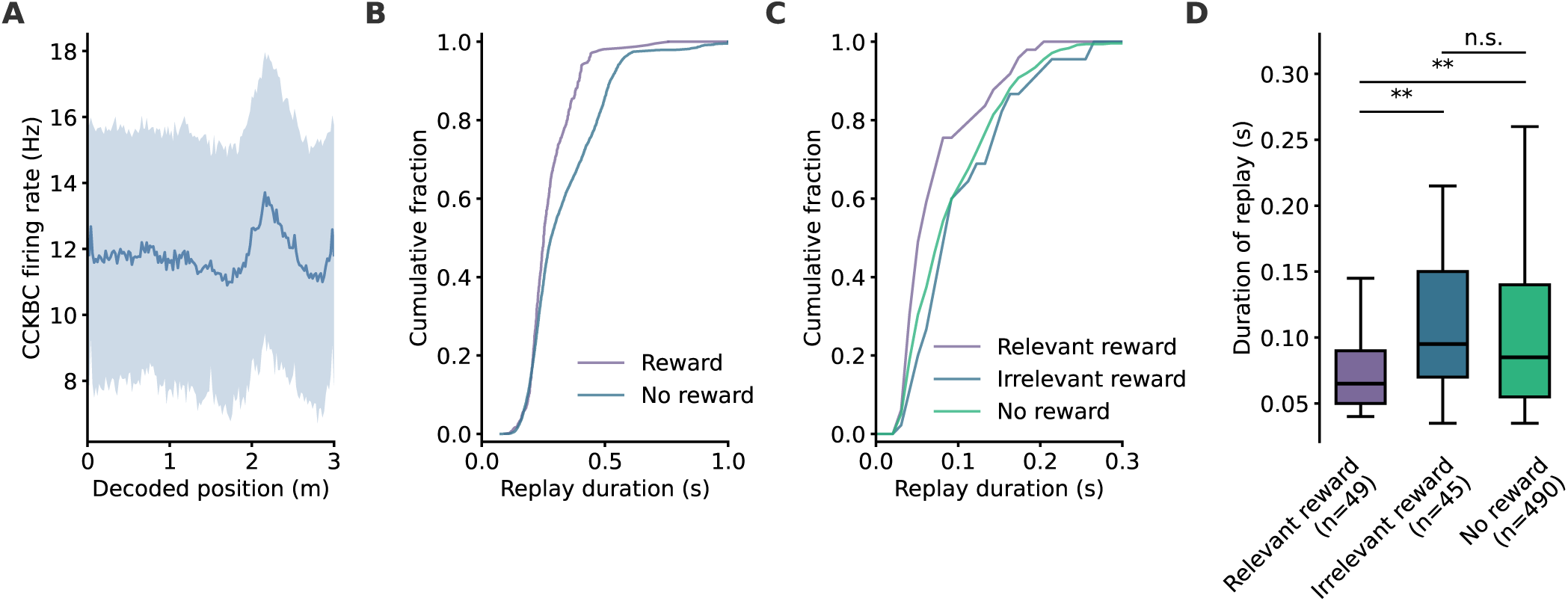
Replays terminating at reward are shorter. ***A***, Firing rate of CCKBCs arranged by decoded position during replay. Shaded areas represent standard deviation. ***B*** , Cumulative distribution of replay durations in the model, sorted by termination site (reward vs elsewhere). ***C*** , Cumulative distribution of replay durations from experimental data, sorted by termination site (reward vs irrelevant reward vs elsewhere). ***D*** , Statistical comparison of replay durations from experimental data, sorted by termination site (reward vs irrelevant reward vs elsewhere). n.s. : not significant, **: p-value *<* 0.01 Data in ***A*** and ***B*** from 500 50-s long simulations.

To verify this behavior in real data, we analyzed neural activity data recorded with a tetrode from the hippocampal CA1 of rodents performing a context-dependent associative memory task that combined navigation in an S-maze and digging for reward at varying locations (data from Chiossi *et al*. 2025). Briefly, we followed the traditional two-step SWR detection method (high activity in both the multi-unit activity and ripple band-pass filtered LFP), then decoded sequences through Bayesian decoding and place field information during exploration, and only kept candidate events with a good linear fit. The reward location was alternated between blocks of 5 consecutive trials within the same context, and we only considered replays occurring during successful trials to maximize the number of replays matching the correct context. See Materials and Methods for details.

This left us with 584 sequence-coding replays, which were sorted into reward-terminating, irrelevant reward-terminating (termination at a location rewarded in another context), and termination elsewhere (no reward). In accordance with our model predictions, we observed a shortened duration for replays terminating exclusively at the contextually relevant rewarded location (Fig. 7C-D, relevant vs. no reward: p-value = 0.004, relevant vs. irrelevant reward: p-value = 0.001, irrelevant vs no reward: p-value = 0.896, one-tailed Mann-Whitney U-test). We note that the duration of irrelevant reward-terminating replays is not significantly different from that of no-reward replays. These results confirm the predictions of our model that replay duration is modulated by reward context and flexibly represents relevant rewards.

## Discussion

We built a computational model of the local CA3 microcircuitry responsible for SWRs and replay. We demonstrated that TORO-to-CCKBC inhibitory synapses critically control the widely ranging duration of SWR events by tuning the timescale of TORO-mediated disinhibition of the PC-PVBC ripple network, and that the recovery of CCKBC activity directly disrupts the ripple oscillation. Our model reproduced the experimentally observed overrepresentation of reward at termination of replay, thus linking replay activity during SWRs with the behaviorally important episodes (i.e., rewards). Somewhat counterintuitively, our model predicted that events terminating at reward-associated locations are shorter than those ending elsewhere, a finding that we verified using experimental data.

### Relation to previous models

Our model builds upon previous computational models of replay (Ecker *et al*. 2022) and SWR dynamics (Schlingloff *et al*. 2014; Evangelista *et al*. 2020). First, our model extends the FINO model of SWR generation (Schlingloff *et al*. (2014)) by incorporating a TORO-CCKBC disinhibition mechanism in bistable dynamics. Fluctuations in the activity of the PC population activate TORO cells, transiently inhibiting CCKBCs and increasing the excitability of the ripple network that comprises PVBCs and PCs. As PC activity increases further, the FINO mechanism recruits PVBCs for ripple phase-locked firing, which in turn promotes ripple phase-locked activity in PCs and triggers a SWR.

Secondly, SWR termination in our model relies on the network mechanism in which a pro- (PVBC) and an anti- (CCKBC) SWR population compete for inhibiting PCs (Evangelista *et al*. 2020). Importantly, these authors introduced synaptic depression from the pro- to the anti-SWR interneurons, such that the latter population receives weakening inhibition during a SWR until its activity returns to a high enough level to terminate the SWR.

Our model clarified how the TORO interneuron subtype regulates the activity of the PVBC-CCKBC subnetworks during SWR through a disinhibitory mechanism. As TORO inhibition on CCKBC weakens, CCKBC activity disturbs the ripple oscillation in an unstable equilibrium until it terminates. We emphasize that TORO cells induce at least two novel features that were absent from the previous models into our microcircuit model. First, SWR duration can be widely varied through the disinhibitory mechanism. Second, and most importantly, the hypothetical reward-distant-dependent PC-to-CCKBC connections make replay termination occur more likely at a reward location, where CCKBCs exhibit stronger excitation. The weak TORO-CCKBC synaptic weight immediately after an event induces a refractory period, during which further events are unlikely to occur.

### Computational and behavioral implications of our model

Our model may represent not only reward locations but also landmarks and locations associated with aversive stimuli through a salience-modulated learning mechanism similar to that described in our results. The increased representation of reward termination in replay is often linked to a denser place field coverage at reward (Hollup *et al*. 2001; Sato *et al*. 2020). However, a more recent study has demonstrated that the two phenomena are not necessarily intertwined (Pfeiffer 2022). Our results agree with this finding and explain how replay termination can overrepresent reward without place field accumulation. How exactly an increased place field coverage at reward affects replay remains an intriguing open question. Given the stochastic nature of replay initiation in our model, we speculate that denser coverage would make reward-associated locations more likely to be picked as initiation locations of reverse replay, rather than termination.

As we demonstrated in our results, we can tweak our model to have different distributions of SWR durations. In particular, it can generate long-tailed distributions in accordance with experimental findings (Fernández-Ruiz *et al*. 2019). This is reminiscent of the perceptual bistability model in which the timescale and amplitude of adaptation (and noise) affect the shape of the lifetime distribution of pseudo-attractor states (Moreno-Bote *et al*. 2007). Our model behaves partly like an attractor model in which a combination of slow processes, such as synaptic depression, adaptation, and noise, induces alternations between irregular firing and the SWRs. Indeed, synaptic depression alone would only generate a narrow gaussian-like distribution of durations, as all events would terminate after the TORO-CCKBC synaptic weight or CCKBC firing rate has reached a certain threshold. In contrast, our results in Fig. 4A-B clearly show that CCKBC firing rate/TORO-CCKBC synaptic weight remains stable for several hundred milliseconds before SWRs terminate. The processes that allow noise amplification to have a macroscopic effect on SWR dynamics may have significant implications in memory consolidation by the hippocampus and other brain regions. However, deciphering these processes is beyond the scope of this work.

### Limitations of our model and open questions

TORO cells were first identified by Szabo *et al*. (2022), where they were found to exist in much of the hippocampus, including the CA1 and CA3 areas. The authors hypothesized that the unique position of TORO cells in the hippocampal microcircuitry places them as ideal candidates for coordinating SWR-associated activity. Importantly, Szabo *et al*. (2022) found that the activity of TORO cells precedes that of PVBC relative to the ripple center. Our model supports these findings and relies on TORO for generation of SWRs by inhibiting CCKBC, thus promoting PVBC activity.

Paradoxically, however, TORO cells were also found to strongly project to PVBC in the hippocampal area CA1 (Szabo *et al*. 2022). We initially simulated a network model with TORO-PVBC projection, but observed no functional differences from simulations without this projection. The TORO-PVBC connection can be thought of as a background hyperpolarizing input, the effect of which we can counterbalance in our model by increasing the excitatory PC-PVBC synaptic weights. Therefore, we decided to exclude the TORO-PVBC connection as it did not contribute to the overall microcircuitry dynamics described in our results. It is also uncertain whether TORO projects to PVBC in the CA3 or if this projection is specific to the local CA1 microcircuitry. Further studies may reveal the computational role of TORO-PVBC connections in memory processing by the hippocampal neural circuits.

We have built our model by using the strong recurrent connectivity of pyramidal cells in CA3, historically described by Li *et al*. (1994). A recent experiment has reported much lower connectivity closer to 1% (Guzman *et al*. 2016), but later findings from several groups put the figure closer to 3-4% (Watson *et al*. 2025) or 9% (Sammons, Vezir, *et al*. 2024; Sammons, Masserini, *et al*. 2025), respectively. Interestingly, these studies also identified subtypes in the PC population (deep vs. superficial, thorny vs. athorny) and showed that recurrent connectivity varies across the different subtypes. A recent study proposed a model of SWRs based on these findings and explored how different subtype connectivity affects the SWRs (Sammons, Masserini, *et al*. 2025). In particular, their model suggested that asymmetric reciprocal projections between thorny and athorny PCs induce a time lag in the activations of these cell populations during SWRs. However, how exactly the activation of the heterogeneous subset of PC subtypes would impact replay activity remains to be clarified.

Sharp-wave ripples and hippocampal replay have long been thought of as inseparable components of memory consolidation. SWRs are the most powerful and reliable promoters for activity replay, and replay in our model also always accompanies SWRs. However, recent evidence has revealed that activity replay can occur in the absence of SWRs (Widloski & Foster 2025). What modifications enable our model to generate replay sequences without SWRs is an intriguing open question.

In conclusion, our model provides a framework through which the recently identified TORO cells coordinate SWR generation by controlling the alternation of PVBC-CCKBC inhibition, suggesting that TORO cells are crucial for initiating SWRs and regulating their duration, but not for their termination. Our model directly links the hippocampal microcircuit with the content of replay in a goal-directed context, with supporting experimental evidence.

## Materials and Methods

We built a computational model of hippocampal area CA3 in the rodent brain in order to study the cellular and network mechanisms behind SWRs and replay. Population sizes were determined based on previous estimates from Ecker *et al*. (2022) for PCs and PVBCs (8000 and 150 respectively) and matched with data from Whissell *et al*. (2015) to reach 160 CCKBCs. Szabo *et al*. (2022) observed that TORO account for about ∼ 0.5% of cells in CA3, which corresponds to 50 cells in our model. The overall size of the network matches that of the smallest volume of hippocampus required for spontaneous SWR generation in vitro (Schlingloff *et al*. 2014).

### Single-cell models

All cells in the network are modeled as conductance-based adaptive exponential integrate-and-fire (AEIF) cells (Brette & Gerstner 2005) following Eq. 1 using the NEST simulator (Spreizer *et al*. 2022).

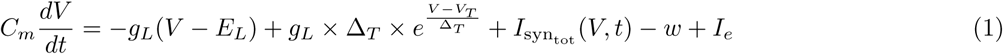

where *C_m_* is the membrane capacitance, *g_L_* the leak conductance, *E_L_* the leak reversal potential, *V_T_* the spike threshold, Δ*_T_* the slope factor of the threshold, and *I_e_* the external input current. The synaptic current *I*_syntot_ is defined as:

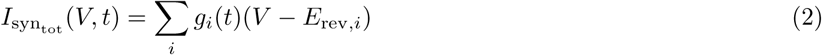

where *g_i_* and *E*_rev*,i*_ are the conductance and reversal potential of the excitatory (AMPA, *E*_rev_ = 0 mV) and inhibitory synapses (GABA, *E*_rev_ = −70 mV) respectively. The spike-adaptation current *w* is defined as:

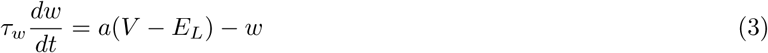

Each time a neuron fires a spike, its adaptation current *w* is increased by a factor *b* and its membrane voltage is reset to *V*_reset_ for a refractory period *t_ref_* . *g_i_*(*t*) in Eq. 2 corresponds to a synapse with biexponential kinetics as in Roth & van Rossum (2009), defined by independent rise and decay time constants (*τ_r_* and *τ_d_* respectively).

Parameters for each modeled population were fitted to experimental data. AEIF parameters for PC and PVBC were taken from Ecker *et al*. (2022) who performed genetic optimization from whole-cell patch-clamp recordings. Following a similar strategy, we obtained recordings from Dudok *et al*. (2021) and Szabo *et al*. (2022) for CCKBC and TORO cells. We extracted membrane capacitance *C_m_*, leak conductance *g_L_*, leak reversal potential *E_L_* and threshold voltage *V_T_* from the whole-cell patch clamp electrophysiological recordings (*E_L_* in TORO was set to −65 mV), and then performed optimization using the NSGA-II algorithm (Deb *et al*. 2002) implemented in the Python package Pymoo (Blank & Deb 2020). The optimization target consisted of equally weighted rheobase, f-I curve, ISI distribution, and ratio of first to last ISI in traces with more than 3 spikes (as described in Dudok *et al*. 2021). Optimization was run for 100 generations with 50 individuals each, and yielded the parameters summarized in Table 1 which were used for all subsequent simulations. Only the parameters marked with an asterisk were obtained through our optimization.

**Table 1:**
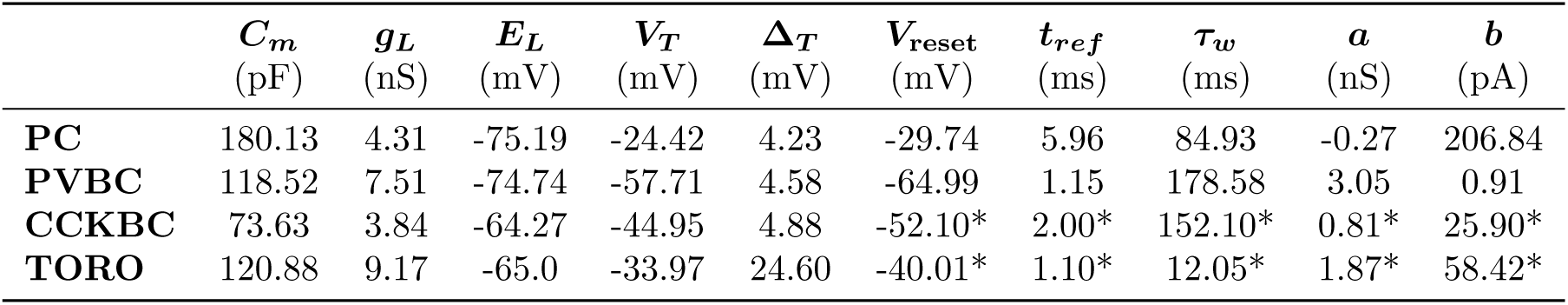
AEIF model parameters for all network populations.

### Network parameters

Network parameters (synaptic strength, short-term depression) were optimized through repeated 10-s long simulations with excitatory input to a subset of 100 PCs with adjacent place fields at *t* = 3.0 and *t* = 7.0 s lasting 0.1 seconds. Parameter combinations (17 parameters in total) were randomly initiated and refined through the NSGA-II algorithm. The following multi-objective function was used to score instances, based on individual population firing rates within and outside of SWRs, ripple frequency in the PVBC population, time, number, and duration of SWRs, absence of gamma power in the PC and PVBC populations:

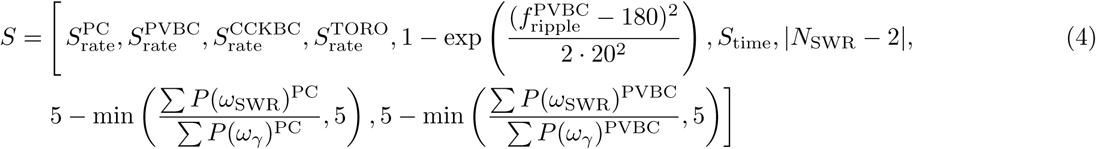

The score for the firing rate of population *p* is given by:

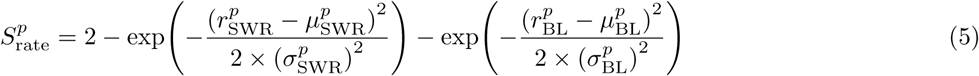

where *r* is the target firing rate during or outside SWRs (*r*_SWR_ and *r*_BL_ respectively), *µ* the observed firing rate, and *σ* the corresponding tolerance. Targets and tolerances for each population can be found in Table 2.

**Table 2:**
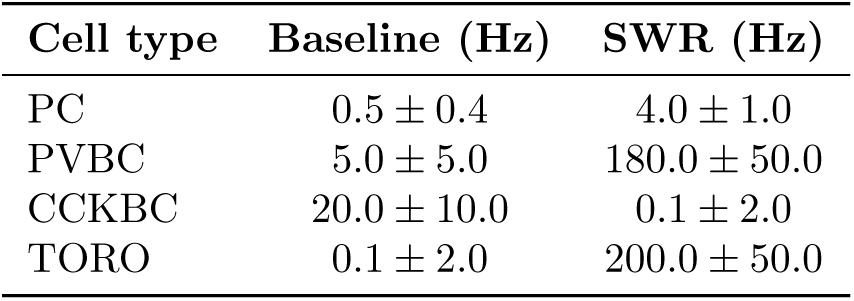
Firing rate targets *r* (± tolerance *σ*) for all cell types at baseline and during SWR for the optimization.

*f* ^PVBC^ in the score for the frequency of the ripple oscillation in the PVBC population is the peak in the 150–220 Hz band determined by Welch’s method. t The PVBC firing rate was segmented into long overlapping windows (500 ms, 50% overlap) tapered with a Hann window to reduce spectral leakage, and their periodograms *P* (*ω*_SWR_) were averaged. Fisher’s g-statistic was used to evaluate the significance of peaks in the ripple band with a p-value of 0.05. In the absence of a significant peak, *S*^PVBC^ was set to 1.

*N*_SWR_ is the number of SWRs observed in the simulation. *S*_time_ scores the timing of SWRs compared to what is expected and is set to 1 if the number of expected vs actual SWRs do not match:

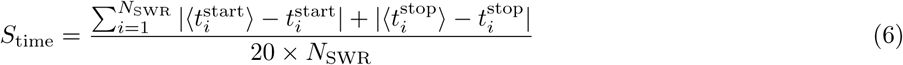

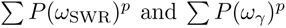 correspond to the total spectral power for population *p* in the ripple (150–220 Hz) and gamma (30–100 Hz) ranges respectively.

After suitable candidates were obtained and tested, excitation at *t* = 3.0 and *t* = 7.0 s was removed and GC-PC weight scaled until SWRs emerged spontaneously (1.1× scaling factor).

All synapse-related parameters can be found in Table 3 along with the optimized weights and connection probabilities. Values are based on data from Ecker *et al*. (2022), Daw *et al*. (2009), Glickfeld & Scanziani (2006), Sammons, Vezir, *et al*. (2024) when available and extrapolated otherwise.

**Table 3:** Connection probabilities, optimized weights, delays, and synaptic time constants between network populations.

| Source | Target | $p_{\text{conn}}$ | $w$ (nS) | Delay (ms) | $\tau_{\text{rise}}$ (ms) | $\tau_{\text{decay}}$ (ms) |
| --- | --- | --- | --- | --- | --- | --- |
| **Excitatory** |  |  |  |  |  |  |
| PC | PC | 0.10 | 0.11–6.18 | 1.2 | 1.3 | 9.5 |
| PC | PVBC | 0.10 | 0.45 | 0.9 | 1.0 | 4.1 |
| PC | CCKBC | 0.10 | 0.7 | 0.9 | 1.0 | 4.1 |
| PC | TORO | 0.25 | 3.93 | 0.9 | 1.0 | 4.1 |
| GC | PC | - | 28.75 | - | 0.65 | 5.4 |
| GC | CCKBC | - | 15.32 | - | 1.0 | 4.1 |
| **Inhibitory** |  |  |  |  |  |  |
| PVBC | PC | 0.25 | 1.68 | 1.1 | 0.3 | 3.3 |
| PVBC | PVBC | 0.25 | 9.05 | 0.6 | 0.25 | 1.2 |
| PVBC | CCKBC | 0.25 | 6.98 | 0.6 | 0.25 | 1.2 |
| CCKBC | PC | 0.20 | 1.74 | 1.6 | 0.9 | 7.3 |
| CCKBC | PVBC | 0.20 | 2.3 | 1.3 | 0.95 | 4.49 |
| CCKBC | CCKBC | 0.05 | 0.54 | 1.3 | 0.95 | 4.49 |
| TORO | PC | 0.05 | 0.27 | 2.0 | 0.85 | 4.6 |
| TORO | CCKBC | 0.20 | 9.86 | 2.0 | 0.85 | 4.6 |

### Short-term depression

We implemented short-term synaptic depression at the TORO to CCKBC synapses according to the phenomenological model of Tsodyks & Markram (1997), using the tsodyks2 synapse model in NEST. For synapses modeled with short-term depression only (*τ*_fac_ = 0), the dynamic utilization variable remains constant (*u_n_* = *U* ). Between presynaptic spikes occurring at times *t_n_* and *t_n_*_+1_, separated by Δ*_t_* = *t_n_*_+1_ − *t_n_*, the fraction of available synaptic resources *x_n_* is updated according to Maass & Markram (2002):

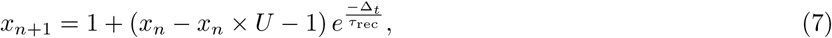

with recovery time constant *τ*_rec_. At each spike, the effective synaptic weight applied to the postsynaptic target is

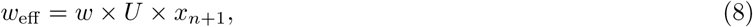

where *w* is the static weight parameter of the synapse described in Table 3. The parameters for short-term depression used in the model are summarized in Table 4. Unless specified, the values used correspond to the default “weak control” implementation described in Fig. 5.

**Table 4:**
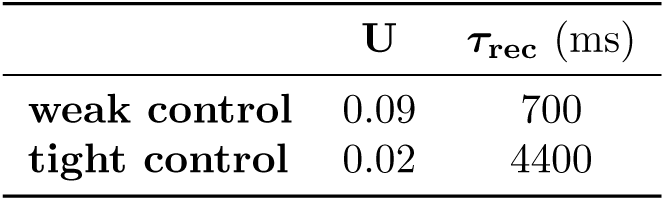
Parameters for short-term depression at TORO-CCKBC synapses.

### Local field potential

Mazzoni *et al*. (2008) showed that the main features of the local field potential can be estimated by a sum of synaptic currents. We implement this as in Ecker *et al*. (2022) by randomly selecting a subset of 400 PCs and summing their synaptic currents then scaling them such that the resulting signal is in the mV range. The obtained LFP has a temporal resolution of 1 kHz. The time-frequency representation in Figure 2B was obtained by calculating the Continuous Wavelet Transform of the LFP with scales spanning frequencies from 1 to 400 Hz using a complex Morlet wavelet with a center frequency of 1.5 and bandwidth of 1.0.

### Place cell identification

#### Modeling

Along the 3-m long simulated periodic environment, place field centers were assigned uniformly to a random subset of 4000 PCs. As place fields are better fit by a thresholded Gaussian characterized by a more abrupt decline with distance (Mainali *et al*. 2025), the tuning curve of cell *i* is defined as follows:

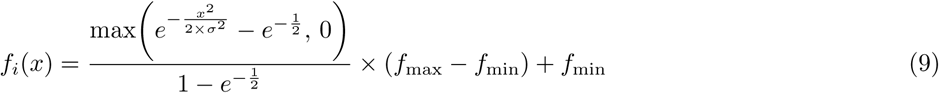

where *x* is the distance to the center of cell *i*’s place field, *σ* = 0.2 m is the width of the place field, *f*_max_ = 15 Hz is the peak firing rate at the center of the place field, and *f*_min_ = 0.1 Hz is the firing rate outside of the place field and matches the baseline firing rate of the remaining 4000 non-place-coding PCs.

#### Experimental

For analyses of experimental replays, place cells are defined from their spiking activity when the speed of the animal (smoothed with a 1 s Gaussian kernel) exceeds 3 cm/s. The tuning curve is obtained from the histogram of spiking activity normalized by occupancy with a 4 cm spatial resolution, smoothed with an 8 cm Gaussian kernel. Only putative excitatory cells (from spike waveform) with spatial information greater than 0.5 bits/s (Skaggs, McNaughton & Gothard 1992) and sparsity smaller than 0.5 (Skaggs, McNaughton, Wilson, *et al*. 1996) were considered. Cells with a peak firing rate below 5 Hz at the center of their place field were discarded. As place cells tend to be orientation-selective especially in 1D environments (McNaughton *et al*. 1983), trials starting from the left and right side were processed separately such that a cell can be place-coding under the criteria above in one direction but not the other.

#### PC-PC weight matrix

The connectivity between pyramidal cells in CA3 reflects the relationship between their receptive fields (van de Ven *et al*. 2016; Sheintuch *et al*. 2023) and previous computational models were able to generate such a connectivity matrix through STDP between spike trains during simulated exploration (Haga & Fukai 2018; Ecker *et al*. 2022; George *et al*. 2023). Importantly, STDP at CA3-CA3 synapses follows a symmetric kernel (Mishra *et al*. 2016), making the learned associations bidirectional. Each PC in the network had a 10% connection probability to any other PC, forming a strongly recurrent network (Li *et al*. 1994; Sammons, Vezir, *et al*. 2024; Sammons, Masserini, *et al*. 2025) with weights following:

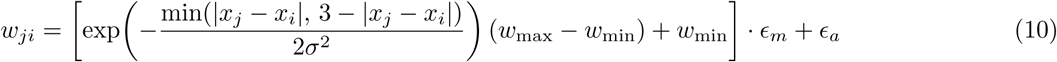

where *x_j_* and *x_i_* are the positions of the place field centers of cells *j* and *i* in the 3-m long periodic environment, *σ* = 0.155 and *w*_max_ and *w*_min_ are the peak in-field and baseline out-of-field firing rates respectively (optimized as network parameters as described above). Multiplicative and additive noise are introduced as:

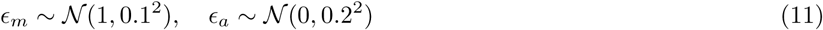

For non-place cells, the distance between *x_j_* and *x_i_* is taken to be the maximal possible distance in the periodic environment (1.5 m). This connectivity scheme approximates that obtained through simulation of spike trains during exploration and STDP as done by Ecker *et al*. (2022) and is less computationally expensive.

#### Replay analysis

In the model, replays are defined as events where the smoothed (50 ms Gaussian kernel) PC population firing rate exceeds 2 Hz for over 0.1 seconds. For experimental data analyses, candidate replay events are defined as periods when the smoothed (10 ms Gaussian kernel) population firing rate recorded at 20 kHz (for spike extraction and clustering, see Chiossi *et al*. 2025) exceeds 3 standard deviations above the mean and extended in either direction to when it crosses the mean again (Pfeiffer & Foster 2013), while the speed of the animal is below 3 cm/s.

SWRs are detected from the processed tetrode recordings (5 kHz) and bandpass-filtered between 150–250 Hz with a 69^th^ order Kaiser window finite impulse response zero-phase shift filter. The absolute value of the Hilbert transform of the signal is applied to obtain its envelope, which is then smoothed with a 50 ms Gaussian kernel to detect events at the sharp-wave timescale. SWRs are defined as events during immobility (speed *<* 3 cm/s) when the signal exceeds 3 standard deviations above the mean and extended in either direction to when it crosses the mean again.

If the detected candidate events in the spiking data did not coincide with the detected SWRs or if the events were shorter than 40 ms, they were discarded from subsequent analyses.

The same replay analysis method was used for both the model and experimental data. We used Bayesian decoding to reconstruct the trajectory from the spikes within a short time window and the tuning curves of the place-coding cells (Zhang *et al*. 1998; Davidson *et al*. 2009). We used log-likelihoods for numerical stability:

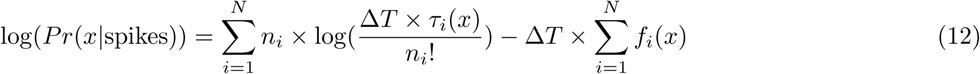

where *N* is the number of place-coding cells, *n_i_*is the number of spikes of the *i*-th neuron in each time window Δ*T* and *f_i_*(*x*) is the firing rate of neuron *i* at position *x* (tuning curve).

To ensure that each time window contains a sufficient number of spikes, we used large overlapping windows (Δ*T* = 20 ms, 15 ms overlap) spanning the whole replay from the first window with at least one spike to the last non-empty window.

Candidate replay events were then scored as in Davidson *et al*. (2009) with a band-finding method assuming constant speed trajectories. For each candidate, we scored goodness-of-fit by a 30-cm wide band with velocities between [-18;18] cm/s (1 cm/s increments) and initial position extending 50 cm in either direction outside the track (6 cm increments). Slow trajectories with speeds between [-5;5] cm/s were excluded as they likely correspond to stationary replay (Denovellis *et al*. 2021). For each event and its best fit speed-initial position pair, 500 shuffles were generated by circularly shuffling the decoded positions by a random amount for each timestep (“column-cycle shuffle” in Davidson *et al*. 2009). Only events with a line fit score over the 99.5^th^ percentile of their shuffled distribution were kept as replay. As each replay is decoded twice (using place fields obtained from either direction walked on the track), only the direction that provided the best fit was kept. The “true” position for each timestep was taken as the weighted mean of the decoded probabilities.

To examine the role of TORO firing rate in SWR generation, candidate SWRs are split into “failed” and “successful”. As the firing rate of TORO is greatly increased during SWRs, classification into “failed” and “successful” SWR-generating signal must be performed prior to SWR start. We found that SWRs in the model were consistently preceded by an increase above 1 Hz in TORO activity (smoothed – 20 ms Gaussian kernel) about 200 ms prior to their start. Any crossing of the 1 Hz threshold was considered a candidate event. Candidates were then split based on whether or not they were followed by a SWR (within 235 ms – empirically determined to match the number of SWRs observed) into either “successful” or a temporary group. The 1 Hz threshold is sensitive to noise and many “non-events” were also flagged into the temporary group. To ensure that only the most relevant candidates were considered and to reduce noise contamination, we only selected the subset of candidates in the temporary group whose sustained activity (in the 50 ms following the threshold crossing) was greater than that of 70% of events in the “successful” group. These events were termed “failed” (as they did not lead to a SWR), and the remaining ones were discarded as falsely flagged.

## Acknowledgments

We thank H. Chiossi and J.Csicsvari for providing us with their experimental data as well as G. Szabo and B. Dudok for making their patch clamp recordings available to us, and S. Fujisawa for sharing his replay analysis code. We are grateful for the support and resources provided by the Scientific Computing and Data Analysis section of Core Facilities at OIST. We thank past and present members of the NCBC lab for their feedback and support throughout the project. This work was supported by JSPS Kakenhi grant JP23H05476.

## Author Contributions

H.M., Conceptualization, Methodology, Software, Formal Analysis, Investigation, Data Curation, Writing – Original Draft, Writing – Review & Editing, Visualization T.F., Conceptualization, Writing – Review & Editing, Supervision, Funding acquisition

The code for this study will be made available on GitHub: https://github.com/orgs/oist-ncbc/repositories

The authors declare that they have no competing interests.

